# REEP6 deficiency impairs ER and Golgi morphologies and causes retinal degeneration by attenuating the expression of phototransduction proteins

**DOI:** 10.1101/2025.03.02.641069

**Authors:** Tongdan Zou, Yongqiong Lin, Ling Li, Yingjie Tong, Ting Wang, Jiajia Wang, Houbin Zhang

## Abstract

REEP6 is a member of the REEP family proteins belonging to the YIP superfamily. REEP proteins were membrane proteins involved in regulating membrane curvature. Mutations in REEP6 are associated with retinitis pigmentosa (RP). Mice lacking REEP6 were generated to model human RP. Reep6 mutant mice exhibit progressive retinal degeneration, which is attributed to abnormal trafficking of PDE6 and the absent expression of guanylate cyclases (GCs), due to ER dysfunction. However, the phenotype regarding the failed expression of GCs is questionable. Here, we generated an independent *Reep6* knockout mouse model. Our *Reep6*-deficient mouse line showed progressive retinal degeneration as previously reported. Nevertheless, unlike the previous report, our *Reep6* mutant mice had normal trafficking of PDE6. GCs were not absent from the mutant mice, but their expression decreased by approximately one third. Other membrane phototransduction proteins, such as rhodopsin and rhodopsin kinase (GRK1) were also reduced. Attenuated expression of membrane proteins might stem from abnormal ER and Golgi function, as in vitro expression of REEP6 altered the expression of the ER marker and Golgi morphologies. Additionally, RNA-seq revealed that deletion of Reep6 caused reduced expression of multiple phototransduction proteins at the transcription level and activated the inflammation pathway. Thus, retinal degeneration associated with REEP6 mutations is the consequence of reduced expression of phototransduction proteins mediated by ER and Golgi dysfunction and ensuing retinal inflammation.

## Introduction

Retinitis pigmentosa (RP) is a group of highly heterogeneous inherited retinal diseases, approximately affecting 1 out of 1/7,000-1/3,000 individuals worldwide (1). To date, more than 70 genes have been associated with non-syndromic RP (https://sph.uth.edu/retnet/). RP mainly affects rod photoreceptors at the beginning of the disease, characterized by night blindness. With the progression of the disease, it also affects cone photoreceptors, leading to complete blindness. *REEP6* (*Receptor Expression Enhancing Protein 6*) has been identified to cause RP in different ethnic groups (2,3). REEP6 is a member of the REEP/YOP1 family belonging to the YIP superfamily (4).

REEP is named so because it was initially shown to enhance the cell surface expression of odorant receptors that belong to the family of GPCRs (G protein-coupled receptors) in cultured cells (5). REEP proteins play an essential role in shaping the endoplasmic reticulum (ER) morphology. Recombinant REEP1 and atlastin are sufficient to organize lipids into the ER-like structure in vitro (6). Mutations in *REEP1* lead to hereditary spastic paraplegia autosomal dominant type 31 (7). REEP1 is also involved in the regulation of intracellular oil droplet formation (8,9). REEP3 and REEP4 determine ER morphologies in metaphase cells during mitosis (10). Thus, different members of the REEP proteins play their unique roles in different tissues.

There are two splicing isoforms of *Reep6* in human and mouse, namely, *Reep6*.*1* and *Reep6*.*2. Reep6*.*1* differs from *Reep6*.*2* by having an additional exon 5, resulting in an additional 27 amino acid residues in the REEP6.1 protein (11). Prenatal mouse retinas exclusively express Reep6.2. The expression of *Reep6*.2 is terminated and replaced by *Reep6*.*1* after the mouse is born. *Reep6*.*1* is only present in rod photoreceptors in the retina. The robust expression of *Reep6*.*1* coincides with the formation of rod outer segments (11), suggesting its important function is specific for rod photoreceptors.

Several *Reep6*-related mouse models have been created to mimic human RP, including knockout and knockin mice (2,3,12,13). Knockout of the Reep6 gene in mice results in progressive degeneration of rod degeneration, recapturing key phenotypes of human RP. Mutant mice with specific deletion of exon 5 only in Reep6.1 exhibit similar phenotypes indistinguishable from those with complete inactivation of *Reep6* (13), indicating *Reep6*.*2* cannot fulfill the function of *Reep6*.*1* in rod photoreceptors.

Ultrastructural analysis of photoreceptors by imaging consecutive sections using electronic microscopy followed by 3-D reconstruction revealed expanded ERs and an increased number of mitochondria in rod photoreceptors lacking REEP6 (2), suggesting that REEP6 is involved in regulating the ER morphology and proliferation of mitochondria. REEP6 is also demonstrated to modulate the expression of multiple proteins in the phototransduction cascade, including rhodopsin. Rhodopsin is markedly downregulated in the retina lacking REEP6. More strikingly, GC1 (guanylate cyclase 1) and GC2 (guanylate cyclase 1) are nearly undetectable in the *Reep6*^-/-^ retina (2). Additionally, PDE6 mistraffics into the inner segments. Based on these results, the retinal degeneration in the *Reep6* mutant was attributed to the absence of GCs (2). However, the requirement of REEP6 for the expression of GCs is questionable. Given that REEP6 is only expressed in rod photoreceptors, and GC1 is expressed in both rods and cones (14), it is unlikely that the deletion of *Reep6* eliminates the expression of GC1 in all photoreceptors including cones that do not express REEP6 in normal mice. Furthermore, GC1 is required for the normal cone function (14), whereas the cone function in *Reep6*^-/-^ is normal. Therefore, retinal degeneration in *Reep6*^-/-^ could be attributed to an unidentified mechanism other than the absence of GCs.

To clarify the elusive mechanism underlying retinal degeneration caused by *REEP6* mutations, in the present study, we generated another independent *Reep6* knockout mouse line. The KO retina exhibited impaired rod dysfunction and progressive retinal degeneration. The expression of rhodopsin is reduced in the mutant photoreceptors. GC1 and GC2 expression is reduced by approximately, but not completely absent. All phototransduction proteins examined, including PDE6, are all correctly targeted to the outer segments. However, the Golgi apparatus is mislocalized to the outer nuclear layer. Ectopic expression of REEP6.1 in cultured cells reduced the level of calnexin, an ER marker, and caused the dispersion of the Golgi apparatus. RNA-seq revealed the activation of the inflammation signaling pathway. Thus, REEP6 is required to maintain normal morphologies of ER and Golgi and ensure the proper expression of phototransduction proteins in photoreceptors Our findings suggest a novel mechanism underlying the mechanism of retinal degeneration associated with REEP6 mutations.

## Materials and Methods

### 1. Animals

The mice were housed in the animal facility at the Sichuan Provincial People’s Hospital. All mice were handled according to the guidelines of the Association for Research in Vision and Ophthalmology for the use of animals in research. All experimental procedures were approved by The Animal Care and Use Committee of the Sichuan Provincial People’s Hospital.

### 2. Generation of the *Reep6* knockout mouse

*Reep6*-knockout mice were generated by using the CRISPR/CAS9 technique. The mouse strain used to generate the mutant mice was C57BL/6J (The Jackson Laboratory, USA). Two gRNAs, gRNA1 (5’GTCTCAAAGAGGAGGAAGAGG3’, matching the forward strand of the gene), and gRNA2 (5’ GGTTCCGATGTTGATGCTGGG 3’, the reverse strand of the gene), were designed to target intron 1 and exon 5 of *Reep6*.*1*, respectively. Deletion of the sequence between the two target sites was screened by PCR using the primer pair F1 (5’ AGACAAGCAGGACTGCCTTG3’) and R2 (5’ CTATACAATCCCTCCCAGAG3’) and verified by sequencing. The mutant allele was genotyped by using the primer pair F1 and R2, which resulted in a 505 bp PCR product. The WT allele was genotyped by using the primer pair F1 and R1 (5’CTGCACTCACGAAGCATATGC3’), yielding a PCR product of 295 bp. The absence of REEP6 in the homozygous knockout mice was verified by immunohistology and western blotting analysis.

### 3. Immunoblotting

Mouse retinas were lysed by brief sonification in 1xSDS loading buffer supplemented with a protease inhibitor cocktail. The proteins in retina lysates were resolved by 10% or 15% SDS-PAGE gels followed by transfer to a nitrocellulose membrane. The following primary antibodies were used at the given dilution: anti-REEP6 (1:2,000, Proteintech, China), anti-GAPDH (1:10,000, Proteintech, China), anti-rhodopsin (1:10,000, custom-made (15)), anti-GC1 (1:5000, Proteinch, China), anti-GRK1 (1:5,000, Proteintech, China), anti-GC2 (1:5,000, a gift from Dr. Wolfgang Baehr at the University of Utah), and anti-PDE6B (1:5,000, Proteintech, China). The secondary antibody used was HRP-conjugated goat anti-rabbit (Proteintech, China) diluted at 1:5,000. The signals were visualized by using an ECL kit (4A Biotech, China). Signal intensities were quantified by ImageJ.

### 4. Immunohistochemistry

Retinal cryosections were prepared as described (16). Retinal cryosections were permeabilized and blocked with 0.2% Triton X-100 supplemented with 5% normal donkey serum (Jackson ImmunoResearch, USA). The sections were immunostained as previously described using primary antibodies against the following proteins: rhodopsin (1:1,000, custom-made (15)), GC1 (1:200, Proteintech, China), GC2 (1:500, a gift from Dr. Wolfgang Baehr at the University of Utah), PDE6B (1:500, Proteintech, China), GRK1 (1:200, Proteintech, China), and GCAP2 (1:500, a gift from Dr. Wolfgang Baehr at the University of Utah). Alexa 488- or Alexa 594-conjugated goat anti-rabbit secondary antibodies (Thermo Fisher, USA) were diluted 1:300. Images were acquired on a Zeiss LSM 800 or LSM 900 confocal microscope (Zeiss, Germany).

### 5. Electroretinography (ERG)

ERGs were performed using the Colordome ERG system (Diagnosys, LLC). The mice were dark-adapted overnight and anesthetized by an intraperitoneal injection (10 ml/kg body weight) of a ketamine (10 mg/ml) and xylazine (1 mg/ml) mixture. The pupils were dilated with 1% tropicamide 10 min prior to the recordings. For scotopic ERGs, the mice were presented with single flashes with varied intensities ranging from 0.002 to 50 cd·s/m^2^. For photopic ERGs, the mice were light-adapted for 10 min under a background light before being presented with single flashes ranging from 1 to 50 cd·s/m^2^.

### 6. Histology

The mouse eyeballs were fixed in 4% paraformaldehyde prepared in 1xPBS at room temperature for 24 hr. The cornea and lens were removed. The eyecup was dehydrated in increasing concentrations of ethanol solutions from 30% ethanol to 100% ethanol. The eyecup was infiltrated with paraffin followed by embedding in paraffin. The paraffin block was sectioned with a thickness of 5 μm. For H&E staining of the section, the section was de-paraffinized by heating the slide at 60 °C for one hour followed by washing with xylene. Subsequently, the section was stained with hematoxylin and eosin (H&E). The row of nuclei in the outer nuclear layer was counted at several different positions across the whole section.

### 7. Construction of REEP6 and rhodopsin expression plasmids

Total RNA was isolated from P30 mouse retinas using TriZol (Thermo Fisher, USA). cDNA was synthesized for by reverse transcription using TransScripi All-in-One SuperMix (Transgen Biotech, China). The mouse full-length Reep6.1 cDNA was cloned into a pEGFP-N1 plasmid (Clontech, USA) with EGFP fused in-frame to the C-terminus of REEP6.1 using NEBuilder (NEB, USA). Similarly, the mouse rhodopsin gene was cloned into pmCherry (Clontech, USA) with mCherry fused to the C-terminus of rhodopsin. The plasmid sequence was verified by Sanger sequencing.

### 8. Cell culture, transfection, and immunochemistry

The COS-7 cell line was obtained from ATCC (ATCC, USA). Culture of the cells followed the standard procedure according to the provider’s instruction. Cells were transfected with the pReep6-eGFP and pRho-mCherry plasmids by using the FuGene transfection reagent (Promega, USA) following the manufacturer’s instruction. Twenty-four hours after transfection, cells were fixed with 4% paraformaldehyde at room temperature for 10 min. Fixed cells were permeabilzed with PBS containing 0.2% Triton-X 100 (Sigma, USA) followed by immunostaining with anti-calnexin (1:100, Proteintech, China) or GM130 (BD Transduction Laboratories, USA). The secondary antibody was Alexa594-conjuated goat-anti-rabbit (1:300, Thermo Fisher, USA) and goat-anti-mouse (1:300, Thermo Fisher, USA). Images were acquired on a Leica confocal microscope (Leica, Germany).

### 9. RNA-seq

Retinas were isolated from WT, *Reep6*^+/-^ (Het), and *Reep6*^-/-^ (KO) mice at the age of one month. RNAs were extracted using TriZol (Thermo Fisher, USA). The RNA integrity was assessed using the RNA Nano 6000 Assay Kit (Agilent, USA). The transcriptome sequencing library was constructed using the NEBNext Ultra RNA Library Prep Kit for Illumina (NEB, USA) following the manufacturer’s instruction. Index codes were added to attribute sequences to each sample. The library preparations were sequenced on an Illumina Hiseq 2500 platform (Illumina, USA), and paired-end reads were generated. These clean reads were then mapped to the reference genome sequence. Differential expression analysis of KO vs wild-type and Het was performed using the edgeR. RNA-seq was performed by Novogene, China.

### 10. Quantitative real-time PCR

Total RNA was isolated from P30 mouse WT and KOretinas using TriZol (Thermo Fisher, USA). cDNA was synthesized described above. Quantitative real-time PCR (qRT-PCR) assays were carried out using the TransStart® Top Green qPCR SuperMix kit (Transgen Biotech, China) with primers specific for genes of interest. *Gapdh* was used as internal control. The primer sequences were listed in **Table 1**. Delta-delta cycle threshold (ΔΔCT = ΔCT (Gene of interest-dCT(Gapdh)) values were calculated and fold change values (FC=2^-ΔΔCT^) were determined.

**Table 1.**
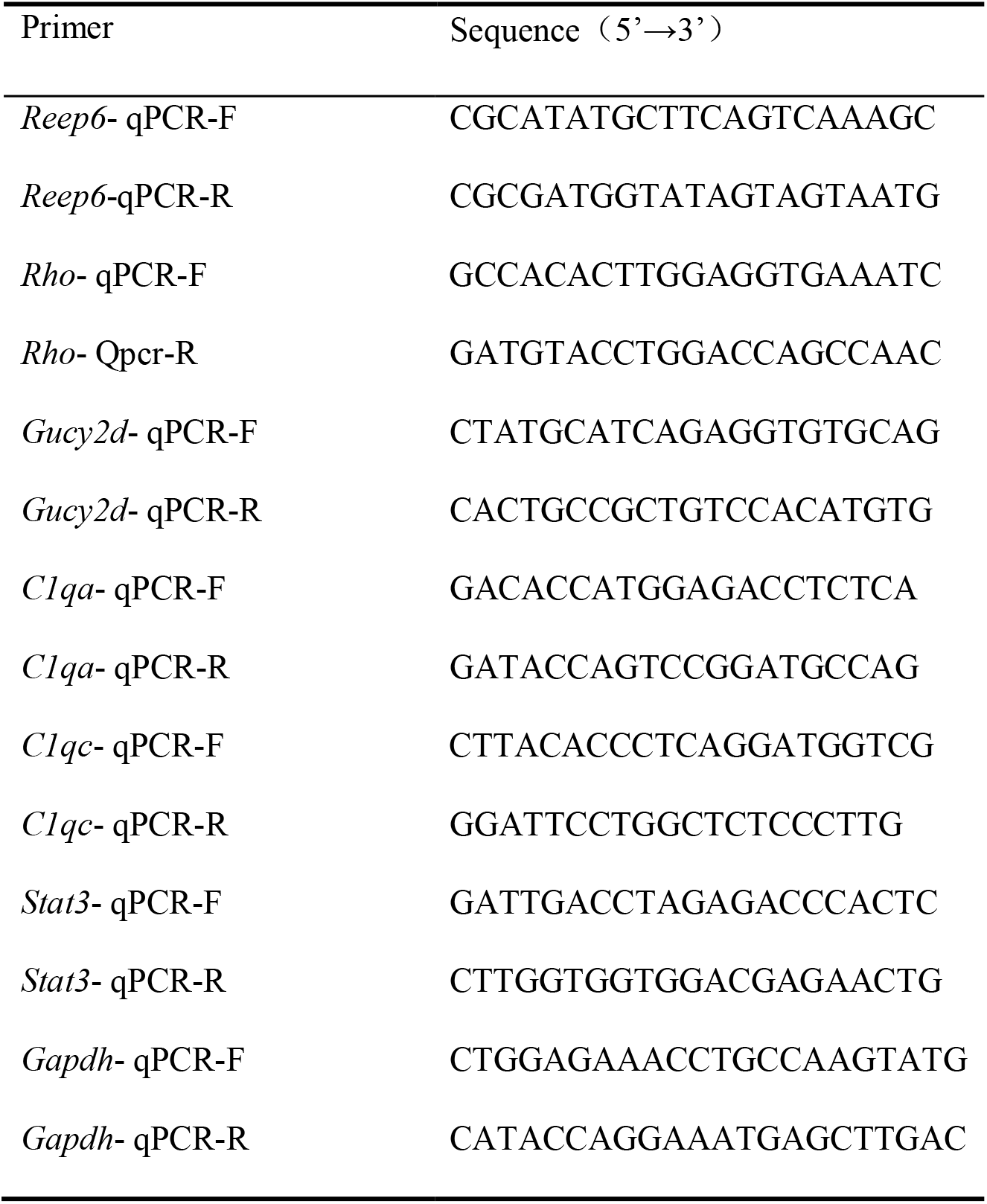
Primers used for qRT-PCR.

## Results

### 1. Generation of the *Reep6* knockout mouse

To explore the in vivo function of *Ree6* in photoreceptors and the pathogenesis of retinal degeneration associated with Reep6, we generated a *Reep6* knockout mouse using CRISPR/CAS9. The knockout strategy is depicted in **Fig. 1A**. Two gRNAs (gRNA1 and gRNA2) were designed to target intron 1 and exon 5, respectively, resulting in the deletion of a sequence encompassing exons 2-4 and part of exon 5. Thus, approximately two-thirds of the coding region was removed, leading to function loss of both REEP6.1 and REEP6.2. Genotyping using PCR confirmed the removal of the sequence between the two targeted sites (**Fig. 1B**). Immunoblotting analysis indicated that there was no REEP6 expression in *Reep6* homozygous knockout (KO) retinas (**Fig. 1C**). As shown by immunostaining, REEP6 was only expressed in wild-type (WT) photoreceptors layers, mainly distributed in inner segments, in agreement with previous reports (2). No REEP6 was detected in the KO retina (**Fig. 1D**), verifying the successful deletion of *Reep6* in the KO. The KO mice appeared normal in development. While the female KO mice bred normally, the male KO mice were sterile, consistent with previous reports (17).

**Figure 1.**
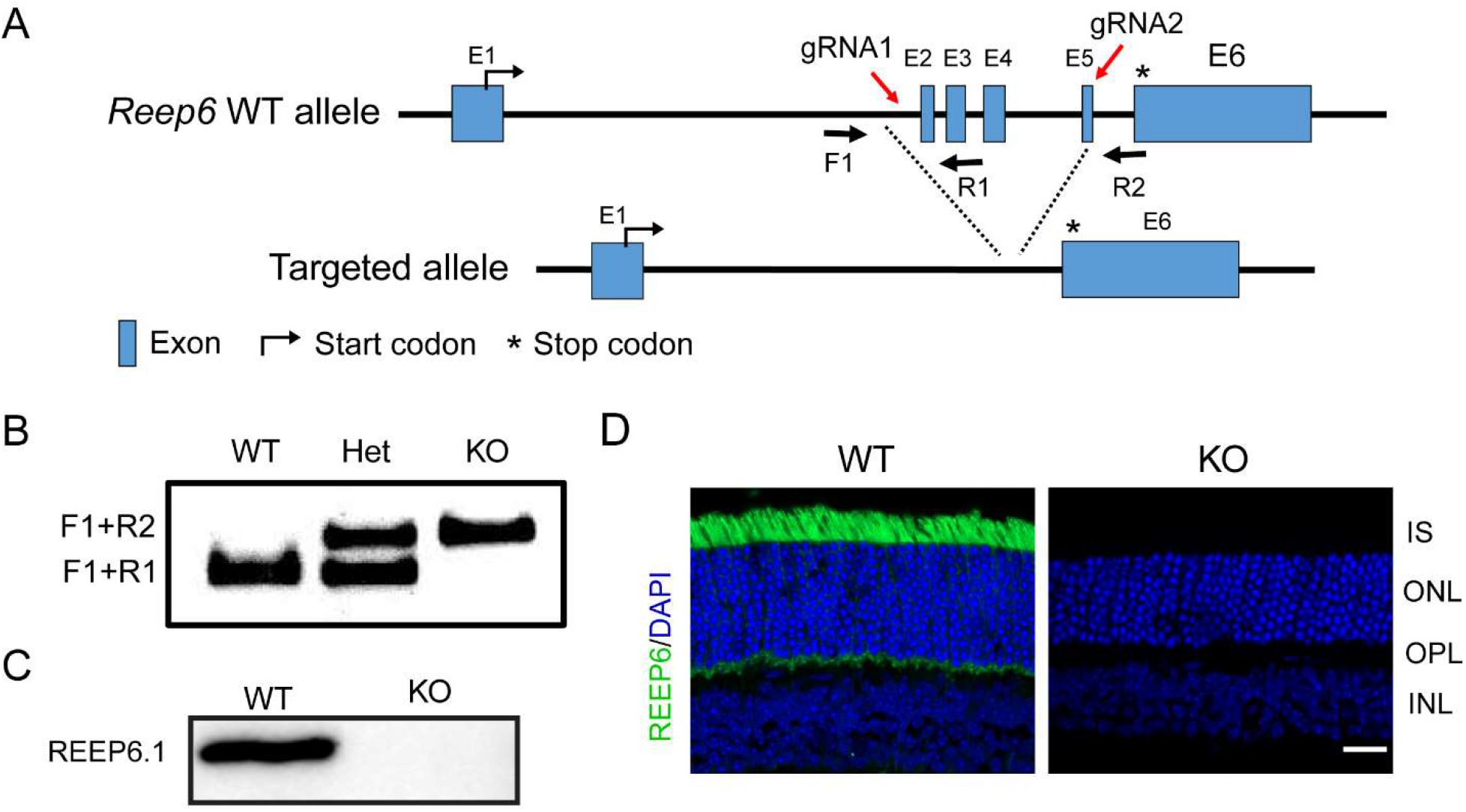
Generation of Reep6 knockout mice. (**A**) Schematic diagram illustrate the knockout strategy using CRISPR/CAS9. Two gRNAs were designed to target intron 1 and exon 5 (E5), respectively. Primers F1 and R2 was used to amplify the wild-type (WT) allele, and primers F1 and R2 were used to amplify the knockout allele. (**B**) Genotyping result for WT, heterozygous knockout (Het), and homozygous knockout mice. (**C**) Immunoblotting analysis to show no REEP6 expression in the KO retina. (**D**) Retinal sections were immunestained with anti-REEP6 (green) and counterstained with DAPI (blue). No REEP6 was detected in the KO retina. IS, inner segment; ONL, outer nuclear layer; OPL, outer plexiform layer; INL, inner plexiform layer. Scale bar, 20 μm

### 2. Progressive retinal dysfunction and retinal degeneration in *Reep6*^-/-^ mice

While *Reep6*.*2* is predominantly expressed in prenatal mice. *Reep6*.*1* is the only *Reep6* isoform that is exclusively expressed in the retina after the mice are born (11). Moreover, *Reep6*.*1* is present only in rod photoreceptors but not in cones. REEP6 deletion is likely to affect rod photoreceptors. To investigate the impact of the REEP6 deficiency on the retinal function, we performed electroretinography (ERG) tests. The scotopic full-field single flash ERGs showed that at the one-month age, the rod function in KO mice was severely affected with decreased a-wave and b-wave amplitudes (**Fig. 2A, B**), whereas the cone function was spared (**Fig. 2C, D**), reflecting the differential expression of REEP6 in rods and cones. At the age of 6 months, while the cone function was slightly reduced in the KO mice, only residual rod function was preserved (**Fig. 2E-G**). At the age of 16 months, rod cells nearly completely lost their function (**Fig. 2G**). In parallel to the progressive loss of rod function, the KO mice also progressively lost their photoreceptors (**Fig 2H, I**). The outer nuclear layer (ONL) thickness in the KO mice was undistinguished from that in the WT control. When the mice reached 6 months old, the ONL thickness of in KO retinas was markedly reduced across the entire retina, and became more severe as the mice aged, indicating progressive retinal degeneration (**Fig. 2H, I**). Thus, Reep6 KO mice largely recapture typical human RP phenotypes, validating the previous reports.

**Figure 2.**
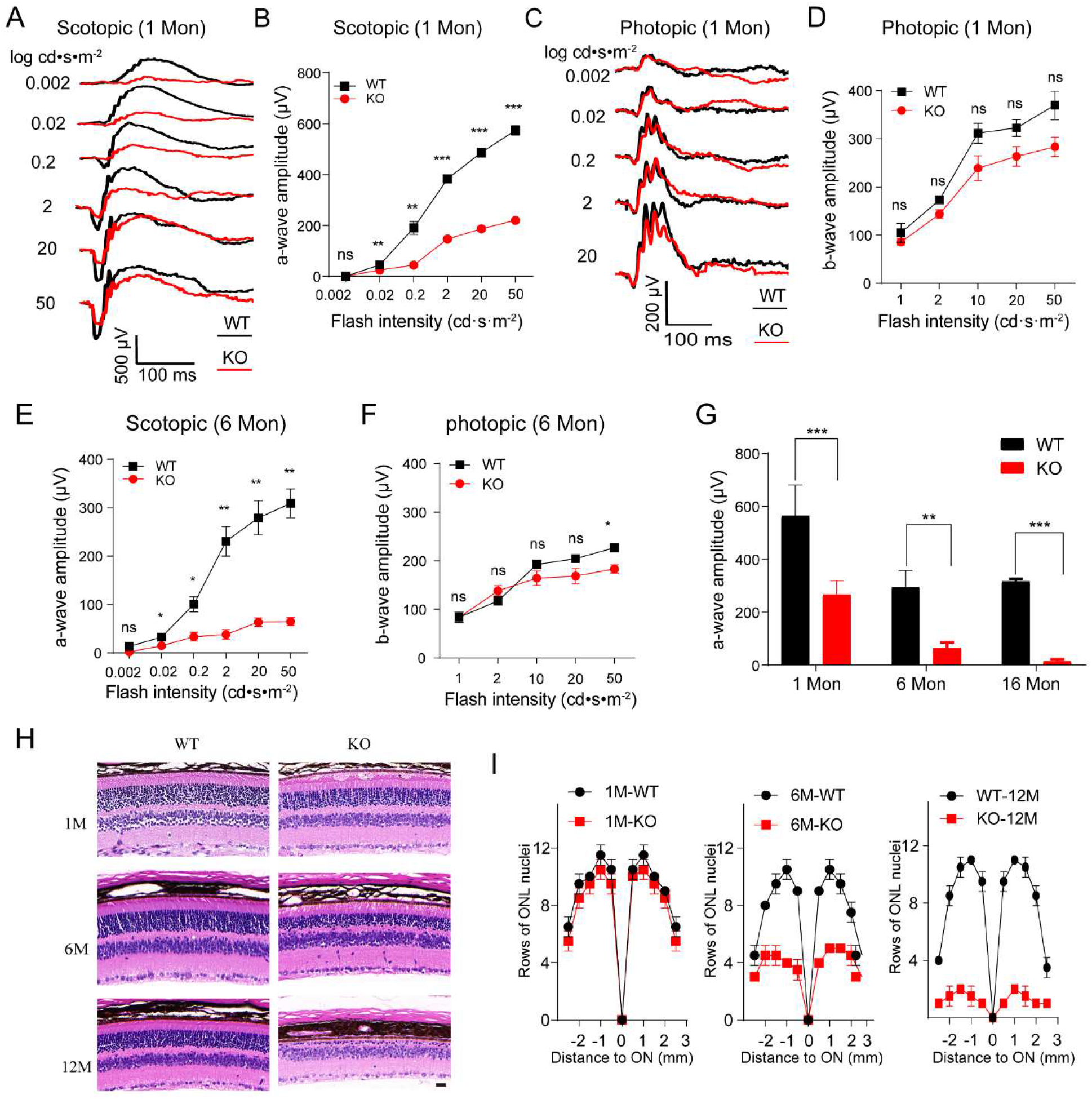
REEP6 deficiency causes retinal dysfunction and progressive retinal degeneration. (**A**) Representative scotopic ERG traces recorded for WT and KO mice at one month of age. (**B, E**) Statistical chart shows a-wave amplitudes of scotopic ERGs recorded for WT and KO mice at the age of one (**B**) and six (**E**) months. (**C**) Representative photopic ERG traces recorded for WT and KO mice at one month of age. (**D, F**) Statistical chart shows b-wave amplitudes of photopic ERGs recorded for mice at the age of one (**D**) and six (**F**) months. (**G**) Comparison of scotopic a-wave amplitudes recorded for WT and KO mice at the age of 1, 6, and 16 months. (**H**) Retina morphology with paraffin-embedded retinal sections stained with hematoxylin and eosin (H&E). Retinal sections were prepared from WT and KO mice at the age of 1 (top panel), 6 months (middle panel), and 12 months (bottom panel). Scale bar, 25 µm. (**I**) Spider graphs depict the total number of rows of nuclei measured at 10 locations of the ONL in H&E-stained retinal sections through the optic nerve. Retinal sections were prepared from WT and KO mice at the age of 1 (left panel), 6 months (middle panel), and 12 months (right panel). n = 6 for each age. Data in this figure were mean ± SD. ns, no significance; \**P* < 0.05, \*\**P* < 0.01; \*\*\**P* < 0.001; unpaired t-test.

### 3. Reduced expression of transmembrane phototransduction proteins

Ectopic expression of a member in the REEP family has been shown to increase the expression of an olfactory receptor, a GPCR protein (5). To assess whether the ablation of Reep6 affects the expression of the most GRCP proteins in rod photoreceptors, we use immunoblotting to analyze the expression of rhodopsin, the most abundant GRCP proteins in rods. As shown in **Fig. 3A, B**, the rhodopsin level was reduced *Reep6*^*-/-*^ retinas at the age of one month, a time point when the number of rods was not noticeably reduced. Immunostaining of retinal sections showed that while rhodopsin was correctly targeted to rod outer segments, rhodopsin was reduced in outer segments of *Reep6*^-/-^ rods (**Fig. 3C**). This result was consistent with the previous report. The expression of other transmembrane phototransduction proteins, such as GC1 and GC2, was also significantly attenuated in *Reep6*^-/-^ retinas (**Fig. 3A, B**). GC1 and GC2 were reduced by approximately 36% and 38%, respectively. Similar to rhodopsin, GCs were correctly localized to photoreceptor outer segments with a slightly reduced amount (**Fig. 3C**). This result is distinct from the previous report showing that GC1 and GC2 were completely absent from the *Reep*6^-/-^ retina (2). Thus, REEP6 deficiency reduces the expression of transmembrane phototransduction proteins, including rhodopsin, GC1, and GC2, but does not affect the transport of phototransduction proteins to outer segments.

**Figure 3.**
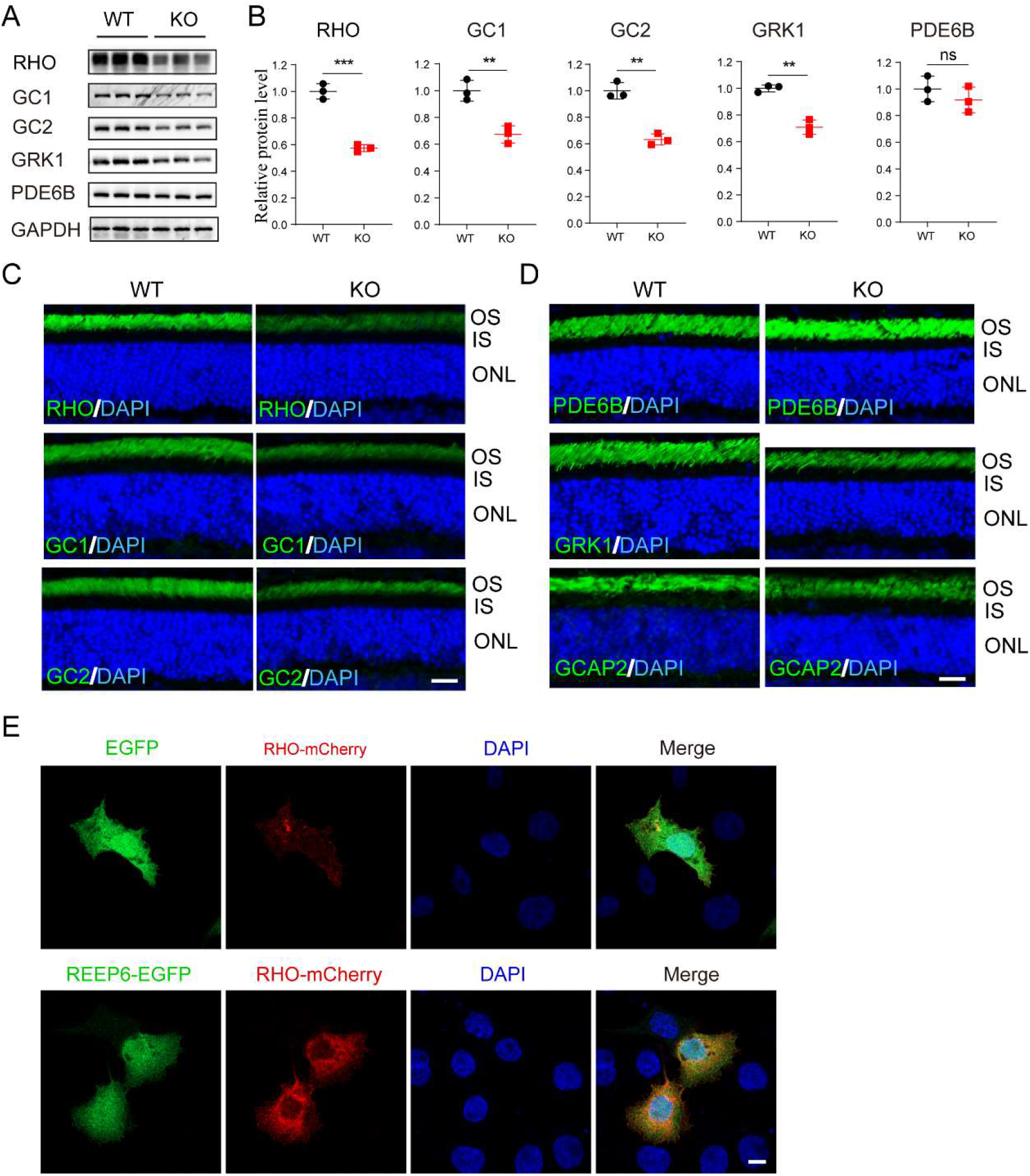
REEP6 promotes expression of phototransduction proteins. (**A**) Immunoblotting analysis of the expression of phototransduction proteins, including RHO, GC1, GC2, GRK1, and PDE6B in retinas of P30 WT and KO mice. GAPDH was used as a loading control. (**B**) Quantification of immunoblotting signals shown in (**A**). Data are expressed as mean ± SD. \**P* < 0.05; \*\**P* < 0.01; \*\*\**P* < 0.001; unpaired *t*-test. (**C, D**) Immunostaining of P30 mouse retinal sections to label RHO, GC1, GC2, PDE6B, GRK1, and GCAP2 (green). Nuclei were label with DAPI (blue). OS, outer segment; IS, inner segment; ONL, outer nuclear layer. Scale bar, 20 μm. (**E**) COS-7 cells transfected to co-express recombinant RHO-mCherry with EGFP (upper panel) or REEP6-EGFP (lower panel). Nuclei were stained with DAPI (blue). Scale bar, 10 μm.

Most phototransduction proteins are transmembrane or membrane-associated proteins. It is not uncommon that the alteration of transmembrane proteins may change the expression of membrane-associated proteins (18). Next, we also examined the expression of membrane-associated phototransduction, such as GRK1, PDE6, and GCAP2. We found that deletion of *Reep6* reduced the expression of GRK1 (**Fig. 3A, B**), but had no effect on its localization of GRK1 (**Fig. 3D**). Similarly, the expression of GCAP2 was decreased in the mutant retina, but its localization was undistinguished from that in WT (**Fig. 3D**). However, the expression or localization of PDE6 was altered, which contrasts with the previous report that showed mislocalization of PDE6 to inner segments. Therefore, *Reep6* deletion affects some membrane-associated phototransduction proteins such as GRK1 and GCAP2, but not PDE6. In addition, *Reep6* is not required for the transport of membrane-associated phototransduction proteins in photoreceptors.

To further validate that REEP6 can promote the expression of transmembrane proteins, recombinant REEP6 was co-expressed with RHO-mCherry in COS-7 cells. Confocal images showed that the expression of RHO was notably increased in cells expressing REEP6 versus mCherry control (**Fig. 3E**). Thus, REEP6 can enhance RHO expression in cultured cells as found in mouse photoreceptors.

### 4. REEP6 modulates the morphologies of the ER and Golgi apparatus

Phototransduction proteins are translated on the ER in photoreceptor inner segments and transported to outer segments via the Golgi apparatus. REEP1, a member of the REEP family, is involved in regulating ER membrane structure. An increase in distal rod inner segment ER volume was previously shown in *Reep6* knockout mice (2), suggesting that REEP6 modulates the ER shaping. To investigate the effect of REEP6 expression on ER, recombinant REEP6 was expressed in COS-7 cells. Staining of the ER to label calnexin, an ER marker protein, indicated diminished calnexin signals in cells expressing REEP6 (**Fig. 4A**), suggesting a reduced ER volume or faster ER turnover. This is in agreement with the finding showing an increase in the ER volume in photoreceptors lacking REEP6.

**Figure 4.**
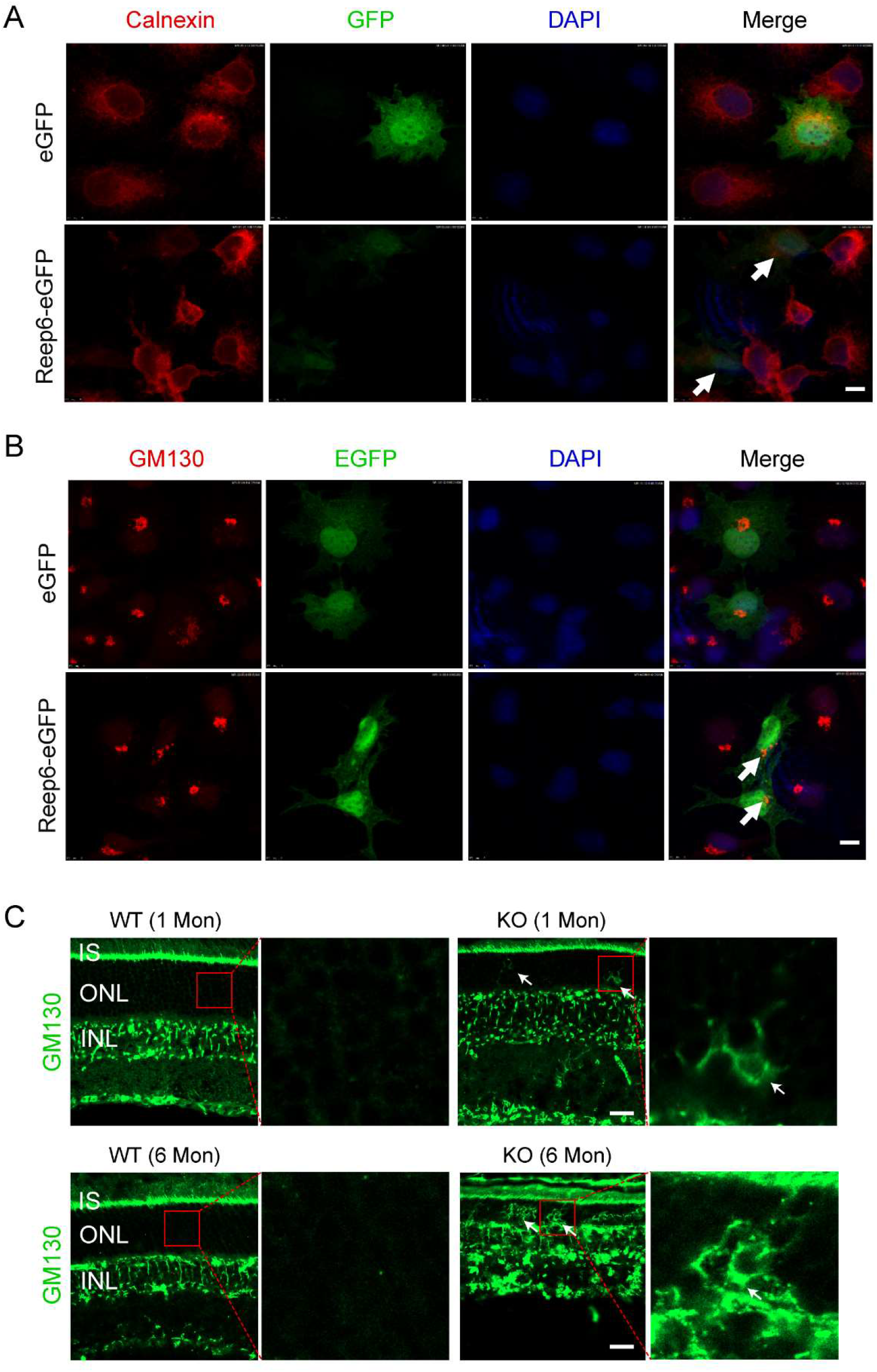
REEP6 modulates ER and Golgi morphologies. (**A**) COS-7 cells were transfected to express EGFP (upper panel) or REEP6-EGFP (lower panel). Cells were stained with anti-calnexin (red) to label the ER. Nuclei were stained with DAPI (blue). Arrows indicate reduced calnexin signal in cells expressing REEP. Scale bar, 10 μm. (**B**) COS-7 cells were transfected to express EGFP (upper panel) or REEP6-EGFP (lower panel). Cells were stained with anti-GM130 (red) to label the Golgi. Nuclei were stained with DAPI (blue). Arrows indicate Golgi dispersal in cells expressing REEP. Scale bar, 10 μm. (**C**) Immunostaining of retinal sections with anti-GM130 (green) to label the Golgi. Retinal sections were prepared from P30 (upper panel) and 6-Mon mice (low panel). Nuclei were label with DAPI (blue). Arrows indicate the Gogli mislocalized in ONLs. IS, inner segment; ONL, outer nuclear layer; INL, inner nuclear layer. Scale bar, 25 μm.

As the Golgi apparatus is part of the membrane system involved in the trafficking of membrane proteins, the effect of REEP6 expression on the Golgi apparatus was also investigated. Compared to the control, REEP6 expression in COS-7 cells resulted in reduced volume of the Golgi (**Fig. 4B**), which was similar to its effect on the ER. Moreover, REEP6 overexpression caused Golgi dispersal. Consistently, abnormal Golgi was also observed in *Reep6* mutant photoreceptors. The Golgi appeared in the outer nuclear layer of the mutant retina, but not in the WT counterpart (**Fig. 4C**).

### 5. REEP6 deficiency causes retinal inflammation

To further gain insight into the mechanism by which retinal degeneration caused by REEP6 mutations, RNA-seq was used to analyze changes in the gene expression profile of retinas from one-month mice lacking REEP6 and controls, including WT and HET (as no apparent phenotype in the HET). Relative to the WT and HET, 527 and 408 genes were significantly upregulated (log2FoldChange > 0.354; *P* < 0.05) and downregulated (|log2FoldChange| < 0.339; *P* < 0.05), respectively (**Fig. 5A**). Examination of genes involving phototransduction in photoreceptors revealed that most of these genes were downregulated, with only one being upregulated, namely, Gnb2 (**Fig. 5B**). KEGG enrichment analysis of differentially expressed genes indicated that other than genes in the phototransduction cascade, several pathways related to virus infection were altered in the mutant retina, such as human CMV infection and HIV infection (**Fig. 5C**), which may suggest the activation of the inflammation pathway. In addition, several gene pathways associated with cholinergic and glutamatergic synapses were changed (**Fig. 5C**), suggesting that *Reep6* may also regulate genes involved in synaptic function. Moreover, several complement factor genes were upregulated in the KO retina, further suggesting inflammation. To verify the RNA-seq results, qRT-PCR was used to quantify the expression of key genes in phototransduction and inflammation. The result showed that *Rho* and *Gucy2d* were downregulated (**Fig. 5D**). In contrast, *C1qa, C1qc, Stat3*, and *Jak3* were upregulated, confirming the activation of inflammation.

**Figure 5.**
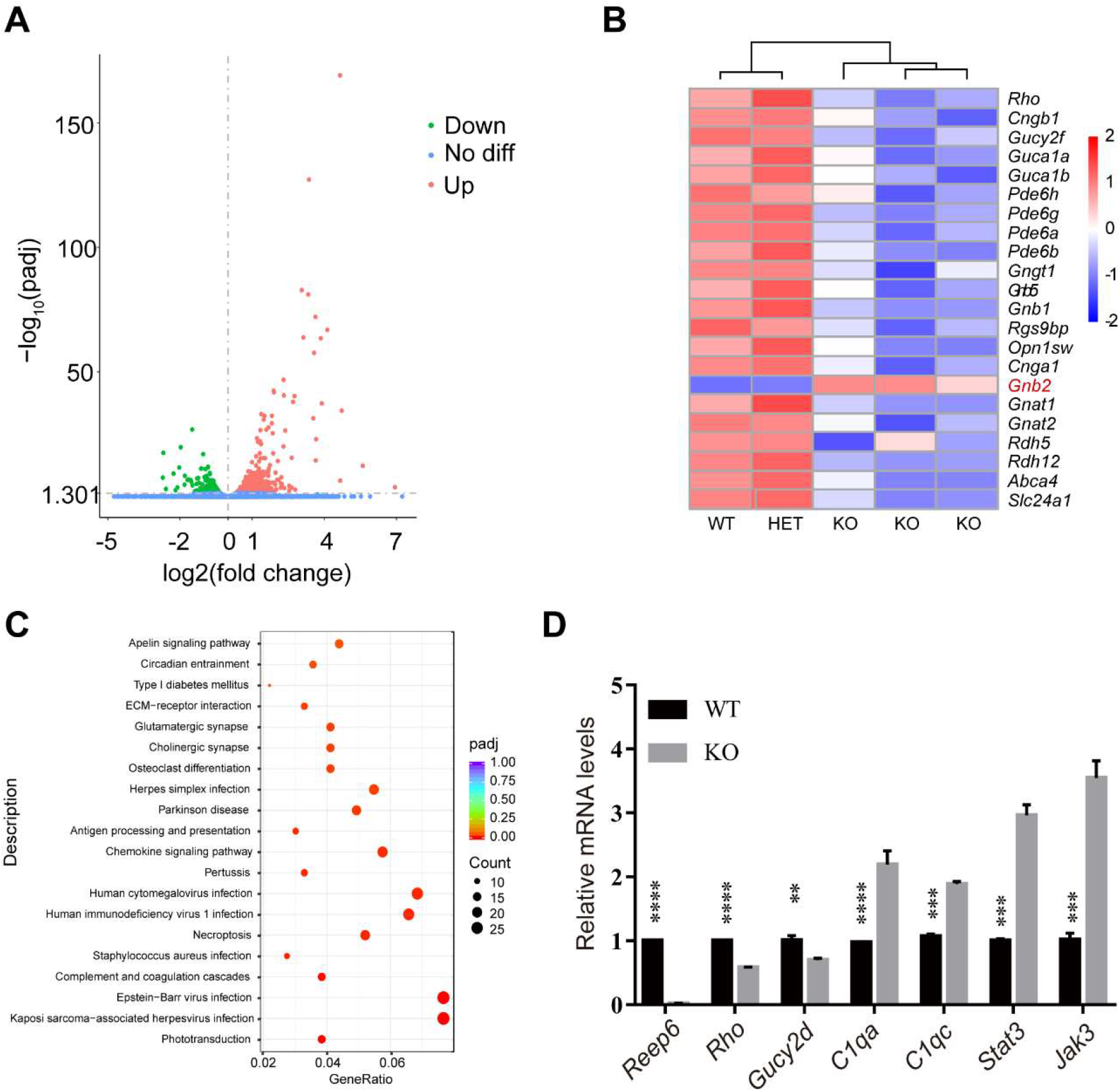
RNA-seq analysis of mouse retinas. **(A)** Volcano plot shows differential gene expression in *Reep6*^-/-^ and control retinas (including WT and HET). (**B**) Heatmap shows differentially expressed genes involved in the rod and cone phototransduction cascades and the retinoid cycles. *Gbn2* is the only gene that is upregulated in KO here. (**C**) KEGG enrichment analysis of differentially expressed genes. (**D**) Validation of the altered expression of *Reep6, Rho, Gucy2d, C1qa, C1ac, Stat3*, and *Jak3* in *Reep6* mutant retinas by qRT-PCR. Data are expressed as mean ± SD. \*\**P* < 0.01; \*\*\**P* < 0.001; \*\*\*\**P* < 0.0001; unpaired *t*-test.

## Discussion

REEP6 has been associated with human RP. Previous studies using mouse models have shown the mechanism could be attributed to defective clathrin-mediated protein trafficking (3), or abnormal mitochondrial fission as well as deficiencies in the expression of phototransduction proteins (2), such as GC1 and PDE6. In this report, we created a new *Reep6* gene knockout mouse. Characterization of the KO mouse showed that the deletion of the Reep6 gene caused rod cell dystrophy and progressive rod retinal degeneration, with cone cells preserved relatively well, which mimics the key features of human RP. Defective energy metabolism and Golgi morphologies were found in the mutant retina.

REEP6 belongs to the REEP protein family. Members in this family are involved in regulating ER morphologies and the maintenance of ER integrities. An in vitro study showed that proteoliposomes containing recombinant atlastin and REEPs form the ER-like tubular network structure (6). This process requires the GTPase activity of atlastin (6). The morphological regulation of ER by REEP1 depends on the hydrophobic domains, which can adopt a hairpin structure and insert into the lipid membrane (19,20). REEP6 shares homologies with REEP1 and contains two conserved hydrophobic domains, which is presumably capable of forming the hairpin-like domain. Therefore, REEP6 is likely able to inset into the lipid membrane and shape the membrane topology. Indeed, the overexpression of REEP6 in culture cells changes the morphology of Golgi. The Golgi apparatus is split into several small portions. Moreover, the overexpression of REEP6 might also change the properties of ER, as the ER marker calnexin was markedly downregulated in cells overexpressing REEP6. According to a previous report, deletion of Reep6 changes ER homeostasis in photoreceptors (2). The ER and Golgi are essential for multiple cellular process in cells. The abnormal ER may severely affect the function of cells. Therefore, REEP6-associated photoreceptor degeneration could be partially due to impaired ER and Golgi functions.

A consequence of the abnormal ER morphology is the change in the number of mitochondria. An increase in the mitochondrial number was documented in the *Reep6* mutant photoreceptors (2). The reason could be that the REEP6 deficiency increases the tubular structures in the ER, which promotes the fission of mitochondria, as the contact between the ER tubules and mitochondria mediates mitochondrial division (21). The mitochondria are the center for energy metabolism. More mitochondrial numbers may suggest more active energy metabolism and produce more ATPs in REEP-deficient photoreceptors. However, the measurement of ATP revealed a decrease in the ATP level in the REEP6 mutant retina (unpublished result). One possible reason for this discrepancy could be that although the mutant photoreceptors have more mitochondria, the mitochondria may not function well, as the protein expression from ER may be compromised. Another factor that could contribute to mitochondrial dysfunction is inflammation, as the RNA-seq revealed that several complement factors were upregulated in the *Reep6* mutant retina, which is a sign of inflammation. Inflammation results in increased ROS production (22), leading to inhibition of mitochondrial function (23). Another possible interpretation is that fatty acids are one of the major fuel sources for mitochondria in photoreceptors. As the ER plays an important role in regulating lipid metabolism, ER stress may affect the lipid metabolism (24), and cut the lipid supply for mitochondria.

Paralleling abnormal ER and Golgi in *Reep6* mutant mice, several proteins involved in rod phototransduction such as rhodopsin and GC2 are also downregulated. Most of the phototransduction transmembrane or peripheral membrane proteins. These proteins undergo posttranslational modification (PTM) in ER and Golgi before transported to the outer segments. Impaired ER and Golgi function may affect PTM and intracellular trafficking of these proteins. Failure to transport these proteins to the outer segment may be degraded by the protein quality control system, thereby reducing the expression of these proteins. Furthermore, a recent study showed that atlastins promote protein exit from the ER (25). Given that atlastins are functionally related to REEP proteins (6), REEP6 may also promote proteins exit from ER or Golgi in phototransduction. Knockout of REEP6 may delay protein exit from ER and Golgi. This could be another factor that causes the downregulation of phototransduction protein in the *Reep6* mutant photoreceptors. Therefore, REEP6 is critical for the efficient expression of phototransduction proteins, particularly in rod cells, as rhodopsin is the most abundant protein in the retina, which accounts for approximately 90% of outer segment proteins (26). Additionally, the rod outer segments renew by disc shedding every 10-15 days (27,28). These exert high demands for protein synthesis and transport in rods. This is in line with the exclusive expression of REEP6 in rods. This hypothesis needs further research to verify.

An earlier report showed no detection of GC1 and GC2 in photoreceptors lacking REEP6 (2), which are different from our findings in this study. If a photoreceptor does not express GCs, it would experience fast degeneration and abnormal development of outer segments (14). Nevertheless, REEP6-deficient photoreceptors degenerate relatively slowly with relatively normal morphologies of outer segments (2), ruling out the absence of GC1 and GC2. GC1 is not only expressed in rod photoreceptors but also strongly in cones and is the only GC involved in the cone phototransduction cascade (14). In contrast, REEP6 is exclusively expressed in rod cells (11). Knockout of *Reep6* is presumed to primarily affect protein expression in rod cells with little in cones at early ages. This rules out the possibility that GC1 is not expressed in the *Reep6*^-/-^ retina. Indeed, our mouse model confirmed that *Reep6*^-/-^ photoreceptors robustly expressed GC1, although modestly downregulated compared to wild-type control. The difference between our result and the reported one likely stemmed from the different procedures used for processing the mouse retinal sections. We used retinal cryosections for immunostaining, whereas the reported study used a paraffin section, which more commonly causes artifacts (29).

In summary, regulating morphologies of ER and Golgi by REEP6 is critical for the normal expression of phototransduction proteins in photoreceptors. REEP6 deficit causes reduced expression of proteins in phototransduction, resulting in retinal inflammation and retinal degeneration.

## Author contributions

T.Z., Y.L, and L.L. designed the experiments, performed the experiments, and analyzed the data. H.Z. conceived the study, designed and supervised the study, performed the experiments, analyzed the data, and wrote the manuscript.

## Funding

This study was supported by the National Natural Science Foundation of China (nos. 82371059 (H.Z.)), 82102470 (J.W.)), and Sichuan Science and Technology Program, China (no. 2024JDRC0048 (L.L.)).

## Data availability

Data are available upon reasonable request.

## Ethics approval

All the procedures involved animals in this study were approved by the Ethics Committee of the Sichuan Provincial People’s Hospital.

## Conflict of Interest

The authors declare no competing interests.

## Notes

### Competing Interest Statement

The authors have declared no competing interest.

### Summary of Updates

1. The name of one of the authors was corrected to "Ling Li" from "Lin Li". 2. Added the funding information of the grant associated with Ling Li.

